# The periodic axon membrane skeleton leads to high-density sodium nanodomains but does not impact action potentials

**DOI:** 10.1101/2021.02.23.432468

**Authors:** Zhaojie Chai, Anastasios V. Tzingonunis, George Lykotrafitis

**Affiliations:** Department of Mechanical Engineering, University of Connecticut, Storrs, Connecticut, USA; Department of Physiology and Neurobiology, University of Connecticut, Storrs, Connecticut, USA; Department of Biomedical Engineering, University of Connecticut, Storrs, Connecticut, USA

**Author notes:** **Corresponding Author** George Lykotrafitis.

## Abstract

Recent work has established that axons have a periodic skeleton structure comprising of azimuthal actin rings connected via longitudinal spectrin tetramer filaments. This structure endows the axon with structural integrity and mechanical stability. Additionally, voltage-gated sodium channels follow the periodicity of the active-spectrin arrangement, spaced ∼190 nm segments apart. The impact of this periodic sodium channel arrangement on the generation and propagation of action potentials is unknown. To address this question, we simulated an action potential using the Hodgkin-Huxley formalism in a cylindrical compartment but instead of using a homogeneous distribution of voltage-gated sodium channels in the membrane, we applied the experimentally determined periodic arrangement. We found that the periodic distribution of voltage-gated sodium channels does not significantly affect the generation or propagation of action potentials, but instead leads to high-density sodium channel nanodomains. This work provides a foundation for future studies investigating the role of the voltage-gated sodium channel periodic arrangement in the axon.

## Introduction

The action potential, a rapid all-or-none change in the membrane potential, is a defining feature of excitable cells like neurons. Hodgkin and Huxley first described the fundamental principles of action potentials 70 years ago, informing generations of scientists on the inner workings of axon currents and their relationship to the generation and propagation of action potentials. In the canonical model, an action potential is triggered when a stimulus in the form of an excitatory event passively reaches the axon initial segment, depolarizing it sufficiently to reach the activation threshold of voltage-gated sodium (*Na*_*v*_) channels [1]. This kindling then leads to an eruption of electrical activity that generates an action potential. The membrane potential returns to its resting state due to a combination of *Na*_*v*_ channel inactivation and delayed activation of voltage-gated potassium channels. In subsequent years, researchers discovered and cloned the multiple sodium and potassium channels that contribute to the action potential as well as the scaffolding and anchoring proteins that organize these channels in the axonal plasma membrane. This work has resulted in a model in which *Na*_*v*_ channels are clustered at high concentrations in the axon through interactions with axonally enriched ankyrin-G and ankyrin-B proteins, which in turn tether *Na*_*v*_channels to the actin-spectrin cytoskeleton.

Although much progress has been made over the last few decades to understand not only the electrical properties of the axon but also its cell biology, it was not until 2013 that the nanoscopic organization of the actin-spectrin skeleton of the axon was revealed [2]. Deviating from previous models, the actin-spectrin skeleton is composed of ring-like F-actin proteins connected longitudinally by αII/βII or αII/βIV-spectrin tetramers. The periodicity of the membrane-associated periodical skeleton (MPS) is ∼190 *nm* set by the spectrin filaments that are under entropic tension [2, 3]. The spatial arrangement of the actin-spectrin proteins differs from the one found in the soma, where the actin-spectrin skeleton is arranged in a two-dimensional polygonal lattice structure [4]. Computational and experimental work has now shown that the MPS bestows the axon with distinct mechanical properties. For instance, atomic force microscopy experiments revealed that the axon is substantially stiffer than the soma and dendrites of hippocampal neurons [5]. Moreover, using a coarse-grain molecular dynamics model for the axon membrane skeleton, Zhang et al. showed that the axonal MPS has two distinct moduli of elasticity that are correlated with its geometric structure, characterized by stiff actin rings connected by extended spectrin filaments oriented along the axon [5, 6]. The role of the MPS in defining the mechanical properties of the axon is also supported by work showing that deletion of spectrin proteins leads to axon deformation and a higher propensity to form kinks and breaks [7, 8]. More recent work has shown that the MPS might restrict axonal protein and lipid diffusion, although the functional implications of these effects on the action potential are not fully understood [6, 9-13].

Despite our understanding of the cell biology of the axonal MPS and its role in setting the mechanical properties of the axon, its role in the action potential has remained unexplored. As mentioned above, the current model of action potential generation and propagation is based on Hodgkin and Huxley’s mathematical description of the action potential [14, 15]. Consequently, multiple investigators have applied the Hodgkin-Huxley (H-H) formalism to understand how sodium and potassium channels control neuronal excitability of different neurons, and to determine the *Na*_*v*_ channel surface density necessary to stimulate an action potential in neurons. However, an inherent assumption of the H-H model is that *Na*_*v*_ channels are uniformly and homogeneously distributed on the axonal plasma membrane. This assumption is inconsistent with recent super-resolution microscopy studies that showed that *Na*_*v*_ channels follow the MPS periodicity and are spaced ∼190 nm segments apart [1]. Therefore, to examine the effects of the *Na*_*v*_ channel periodicity on the action potential, we simulated the action potential in a cylindrical compartment using the H-H model with the *Na*_*v*_ channels following the experimentally established periodic arrangement rather than being evenly distributed in the membrane.

## Method

Here, we discuss the methodology we followed to solve the H-H equation numerically:

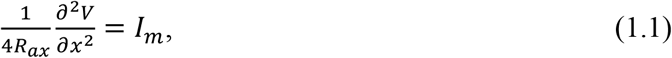

where *Rax* is the axon core resistivity, *V* is the membrane potential measured from its resting value, and *I*_*m*_ is the membrane current density [14, 16].

In the active cable model, the membrane current density is the sum of the potassium current *I*_*K*_(*x, t*), the sodium current *I*_*Na*_(*x, t*), the leak current *I*_*L*_(*x, t*), and the capacitive current *I*_*C*_(*x, t*).

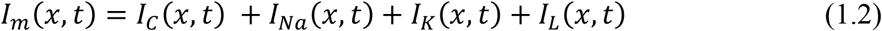

The currents are given by the expressions:

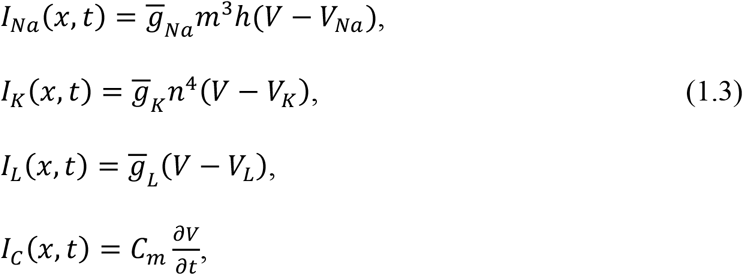

where 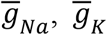, and 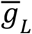 are the maximum sodium, maximum potassium, and maximum leak conductances, respectively (Table 1). *V*_*Na*_, *V*_*K*_, and *V*_*L*_ are the sodium, potassium, and leak reversal potentials, respectively (Table 1). *m*(*x, t*), *h*(*x, t*), and *n*(*x, t*) are dimensionless variables that are used to describe the kinetics of voltage-dependent ionic currents and obey the following equations:

**Table 1:**
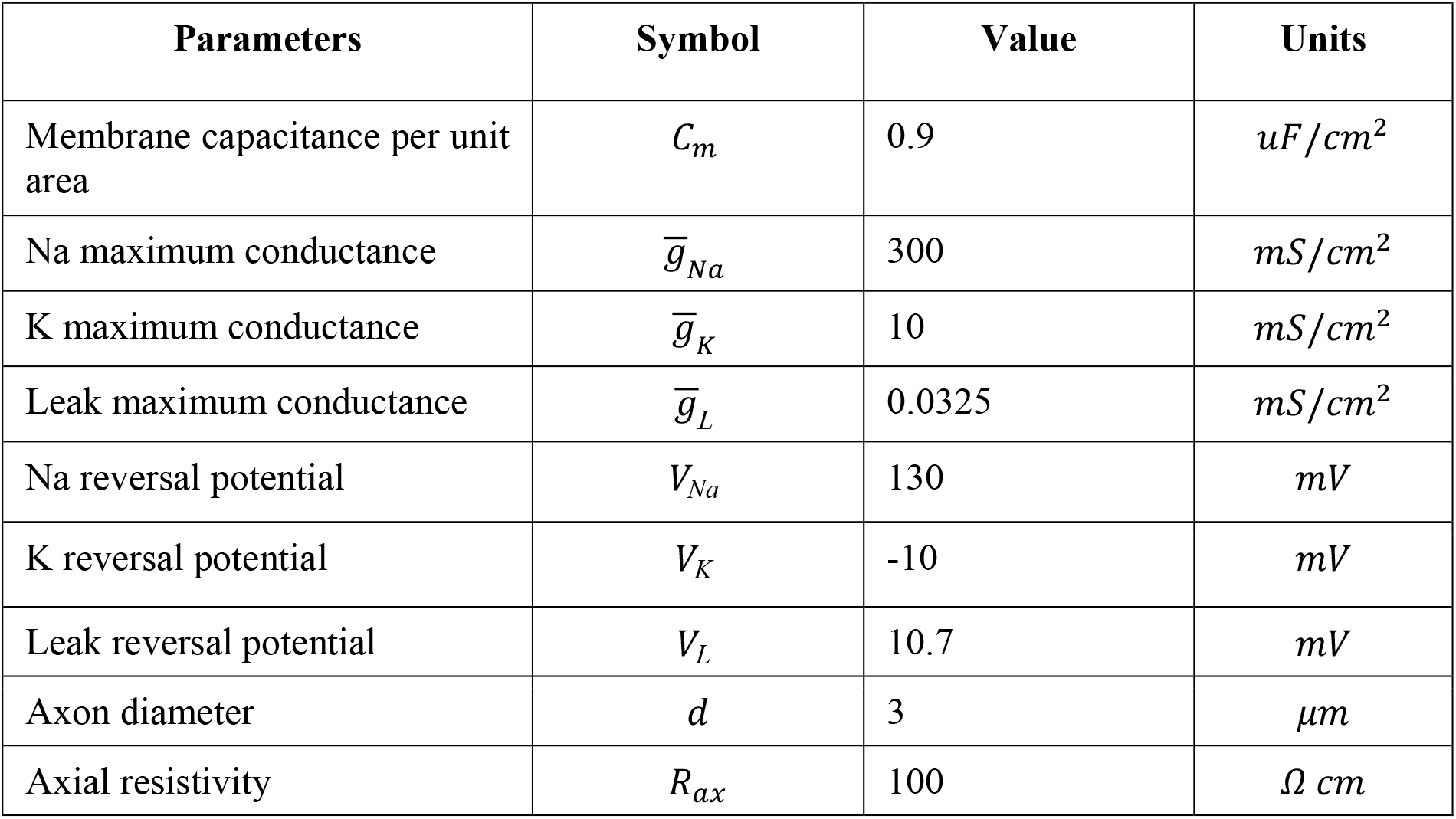
The parameters used in the Hodgkin-Huxley model.

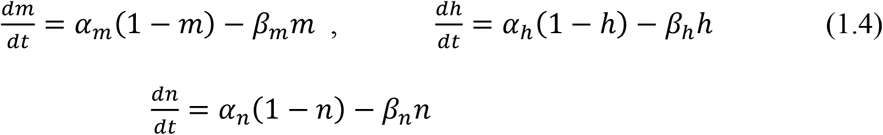

The rate functions that appear in the first-order kinetic equations 1.4 are functions of the membrane potential and are given by the expressions:

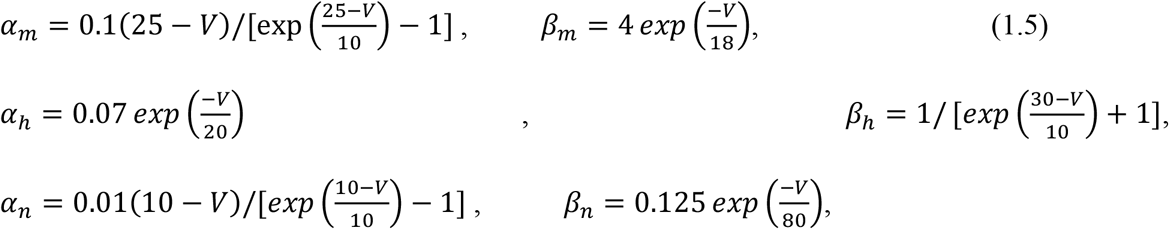

Hodgkin and Huxley obtained the expressions of the rate coefficients by fitting data from several experiments in equations 1.4 [14, 17].

We used the Neumann boundary conditions for the initial-boundary value problem of the H-H equation [18], which correspond to injecting a variable current at one end of an axon obeying H-H kinetics:

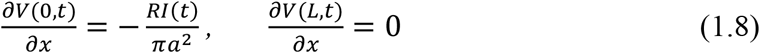

where *I*(*t*) is the injected current at *x* = 0.

The initial conditions are the following:

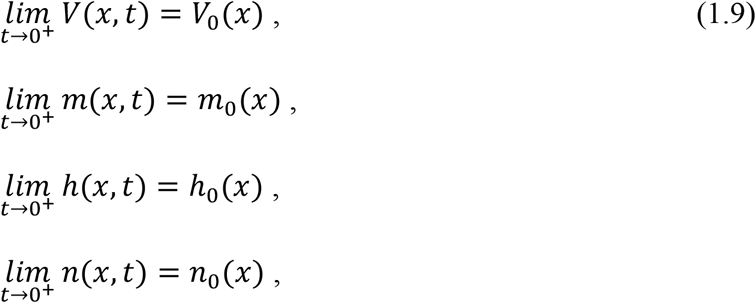

Hence, given the above initial conditions for the rate variables and the potential, we can calculate the values of the rate variables *m*(*x, t*), *h*(*x, t*), and *n*(*x, t*) as a function of time.

To solve the H-H system of the equations discussed above using the initial and boundary conditions above, we employed the backward Euler scheme in MATLAB [19, 20]. We defined a computational grid with time step ∆*t* and number of spatial grid points *N*. The spatial discretization along the axon has a uniform unit length ∆*x* = *L*/*N*. The grid contained *N* + 1 points numbered from zero (corresponding to *x* = 0) to *N* (corresponding to *x* = *L*).

As described in the introduction, a *Na*_*V*_channel can bind to ankyrin, which is tethered to the 15^th^ repeat of β-spectrin near its carboxyl terminus located in the middle area of a spectrin tetramer [21]. This means that a *Na*_*V*_ channel connected to ankyrin is also located in the middle area of two consecutive actin rings. Spectrin tetramers that are not under tension have an end-to-end equilibrium distance of 75 *nm*. In the axon, however, spectrin tetramers are under tension with a reduced range of thermal motion. After considering the size of actin particles, researchers have determined the end-to-end distance of spectrin tetramers in axons to be approximately 150 *nm* [5,22]. The mean distance between two consecutive ankyrin proteins along the longitudinal direction when the spectrin is under tension is *L*_*t*_ = 185.78 *nm* [5]. Because the spectrin tetramers are under entropic tension, the trajectory of ankyrin particles describes a small area around the equilibrium position; consequently, the connected *Na*_*v*_ channels maintain an ordered configuration as they are distributed in azimuthal bands between the actin rings. [2, 5, 6]. Work has shown that the distance of an ankyrin particle from its mean position during thermal motion along the longitudinal direction follows a Gaussian distribution with a standard deviation σ = 0.027 *L*_*t*_, where *L*_*t*_ = 185.78 *nm* in the actual axon (Fig. 1C) [5]. We considered that the width of the band within which the *Na*_*v*_ channels are distributed is 0.108 *L*_*t*_, which is four times the standard deviation of the ankyrin particles and almost 1/9 of the distance between two consecutive actin rings.

**Fig 1.**
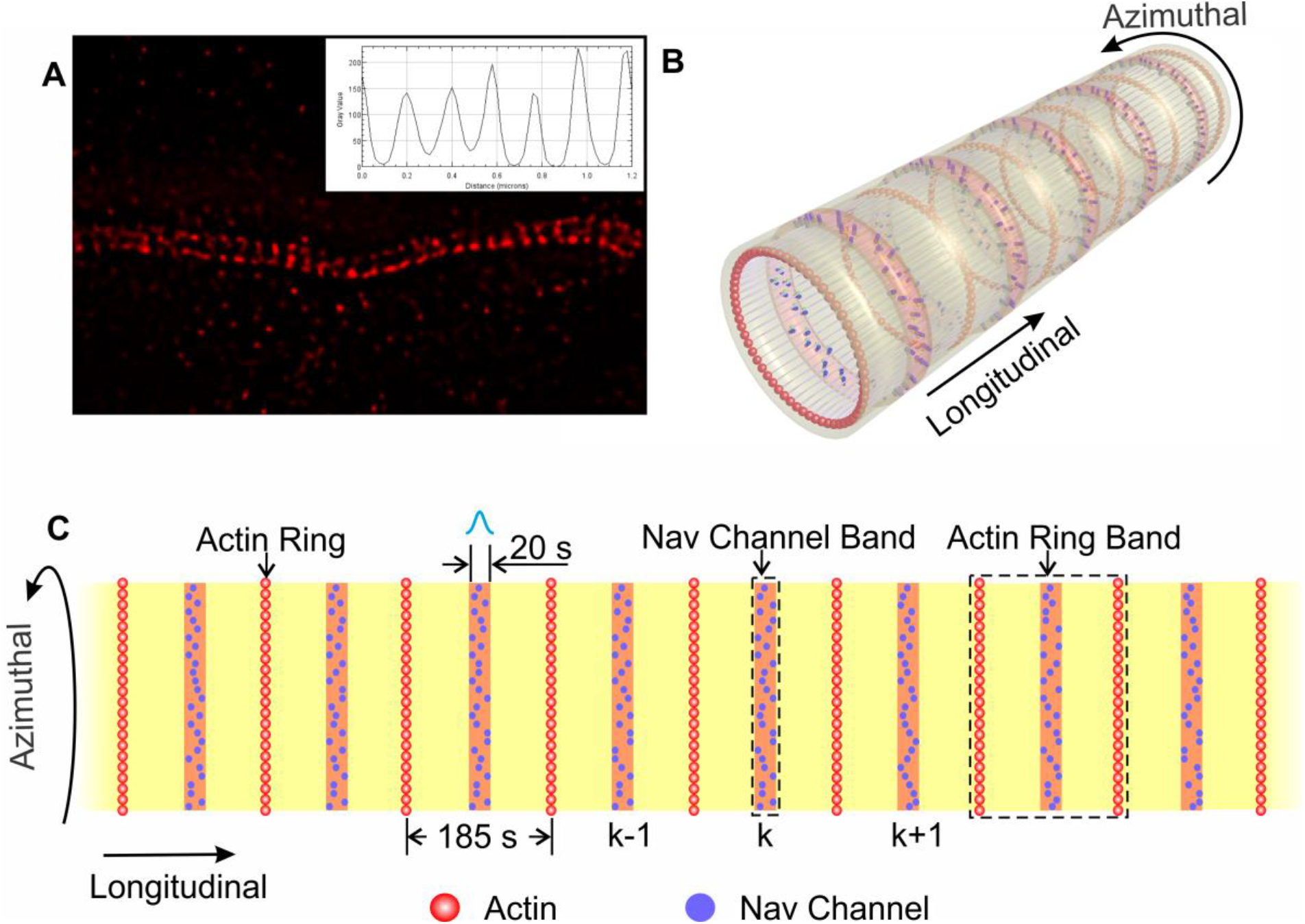
Periodic Gaussian distribution of *Na*_*V*_ channels on the axon. (A) STED image of a single axon recorded from a wild-type (WT) mouse shows the periodicity of the distribution of *Na*_*V*_ channels along an axon. The insert plot shows the intensity profile of the fluorescent image along a line in (A). (B) Illustration of an axon membrane skeleton model based on super-resolution microscopy results. (C) Two-dimensional illustration of the periodic Gaussian distribution of *Na*_*V*_ channels on the axonal plasma membrane considered in this paper. The *Na*_*V*_ channel band width is 20s, approximately 1/9 of the equilibrium distance between consecutive actin rings.

To investigate the effect of the periodic distribution of *Na*_*v*_ channels on the action potential compared to the conventional assumption of homogeneously distributed *Na*_*v*_channels, we implemented two types of *Na*_*V*_ channel conductances *g*_*Na*_ in our simulation. In the first case, we assumed that *Na*_*v*_ channels are distributed homogeneously along the axon. Thus, we adopted a homogeneous *Na*_*V*_ channel conductance 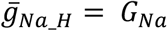, where *G*_*Na*_ = 300 *mS/cm*^2^. In the second case, we considered that the location of *Na*_*V*_ channels follows a Gaussian distribution in bands located in the middle areas between actin rings with a width of 1/9 of the distance between two consecutive actin rings, as mentioned above. It is extremely computationally expensive to use the actual scale *L*_*t*_ = 185.78 *nm* for the numerical solution of the H-H equations. Instead, we adopted a much larger distance between consecutive actin rings and maintained the proportions of the width of the *Na*_*v*_ channels’ azimuthal band with respect to the distance between consecutive actin rings. In particular, we chose 20*s* as the width of the *Na*_*V*_ channel band and 185*s* as the distance between two consecutive actin rings. Then, we chose *s* = *k e*, where *e* is the length scale and *k* is a parameter, and showed that the numerical solutions converge as *k* increases. We obtained plots (Fig. 2 and Fig. 3) for *k* = 20 and *e* = 4.45 *nm*. Consequently, the maximum conductance 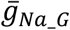 for periodically distributed *Na*_*v*_ channels is given by the expression:

**Fig 2.**
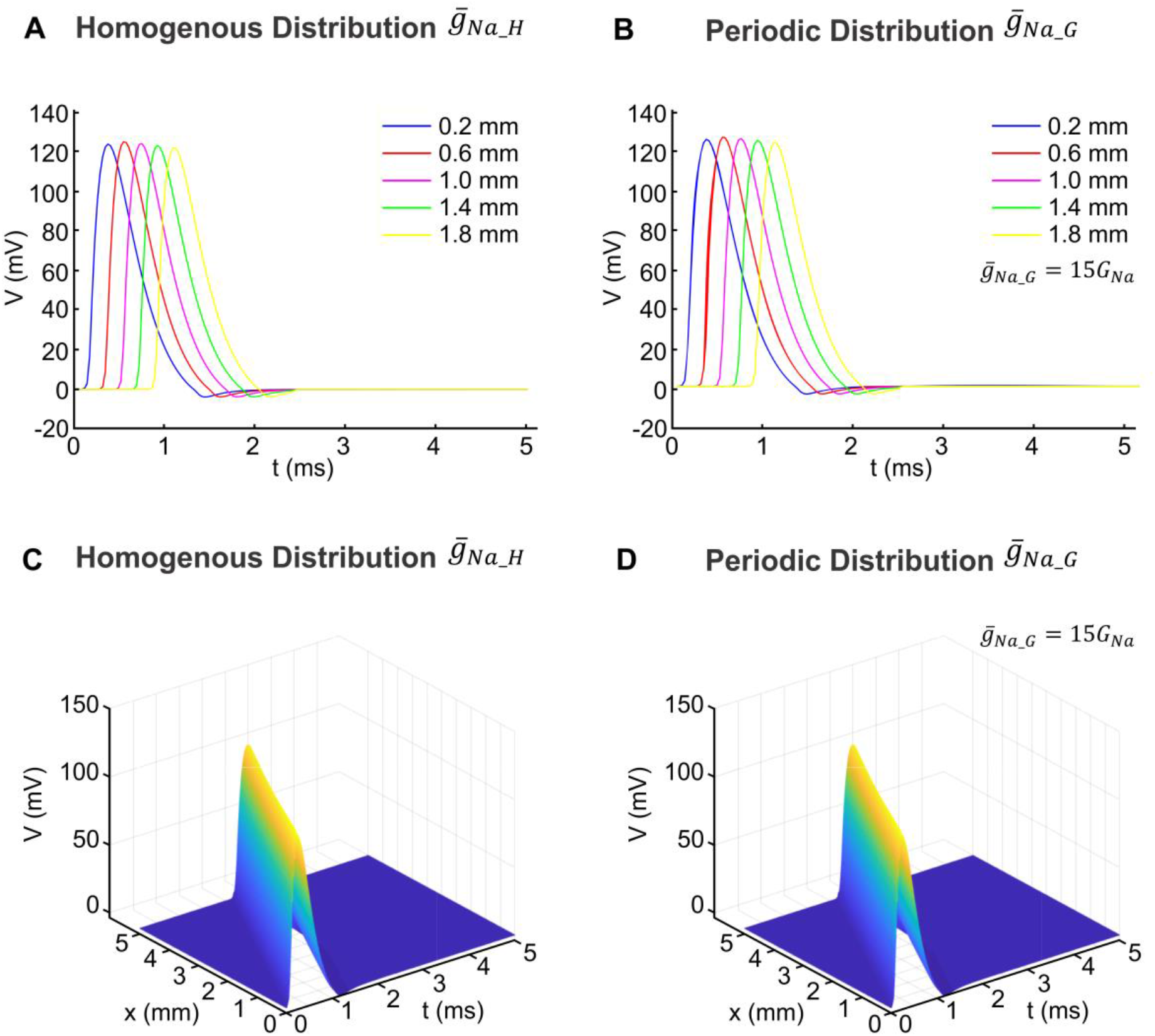
Illustration of action potential propagation for homogeneous and periodic *Na*_*v*_channel distribution. The resting membrane potential was 0 *mV*. (A) Illustration of action potential propagation for homogeneously distributed *Na*_*V*_ channels. The homogeneous *Na*_*V*_ channel conductance was 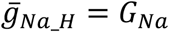, where *G*_*Na*_ = 300 *m*S/c*m*^2^. (B) Illustration of action potential propagation for periodically distributed *Na*_*v*_ channels with a conductance 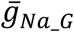 following the Gaussian distribution of Eq. 1.10. (C) Illustration of three-dimensional (3D) action potential propagation along the axon and time for homogeneously distributed *Na*_*V*_channels and conductance 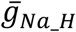. (D) Illustration of 3D action potential propagation along the axon and time for periodically distributed *Na*_*V*_ channels and conductance 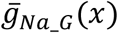.

**Fig 3.**
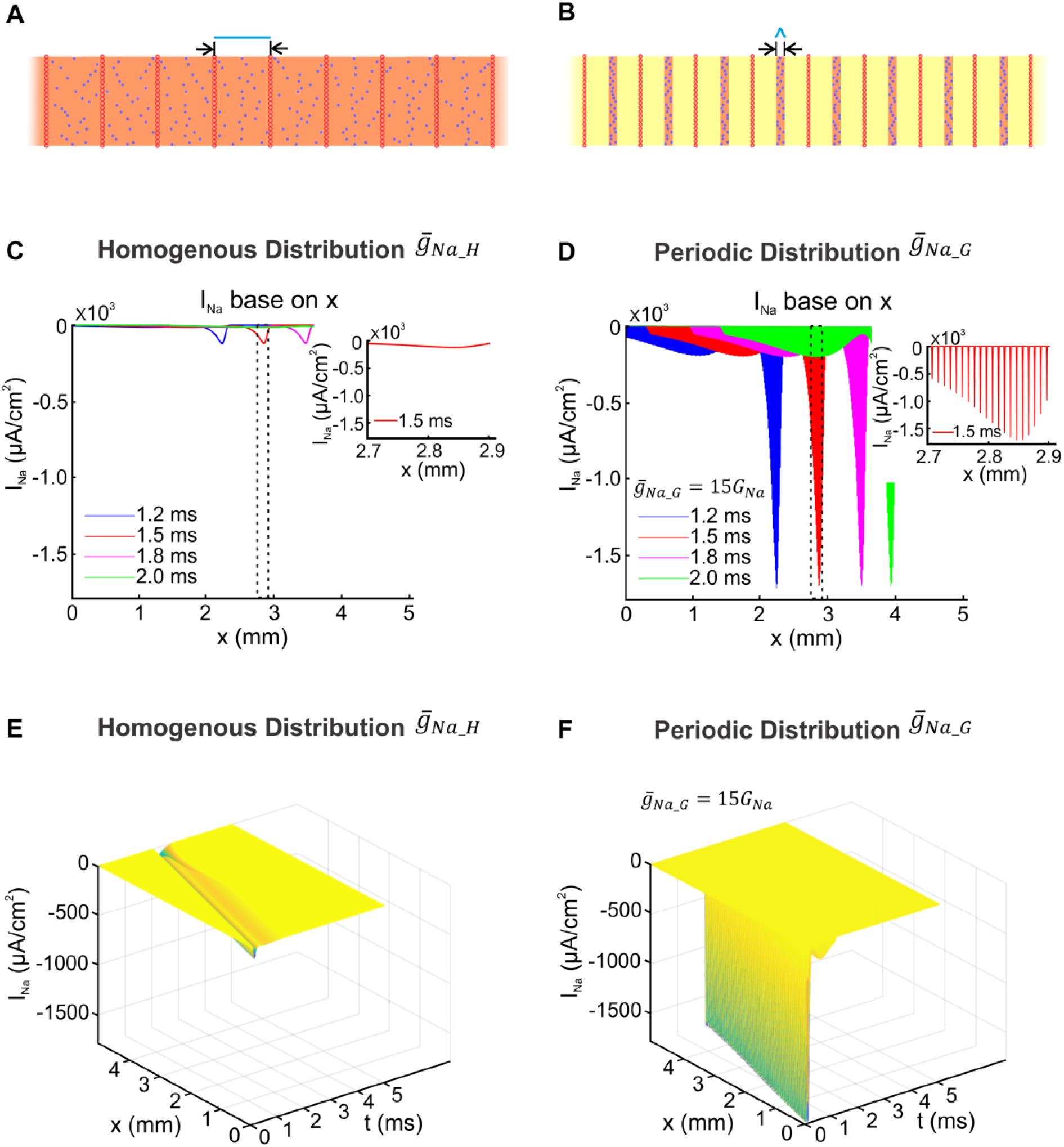
Illustration of the sodium current propagation for homogeneous and periodic *Na*_*v*_ channel distribution. (A-B) Illustrations of the axonal plasma membrane with (A) homogeneously or (B) periodically distributed *Na*_*V*_ channels. (C-D) Plots of sodium current density propagation for (C) homogeneously or (D) periodically distributed *Na*_*v*_ channels. The insert figure is a magnification of the peak of the sodium current where the sawtooth shape of the current is clearly visible. (E-F) 3D plots of the sodium current density along the axon and time for (E) homogeneously or (F) periodically distributed *Na*_*v*_channels. Blue, red, magenta, green, and yellow lines in (C) and (D) are the sodium current densities as function of the position *x* at 1.2 *ms*, 1.5 *ms*, 1.8 *ms*, and 2.0 *ms*, respectively, from the beginning of the action potential propagation.

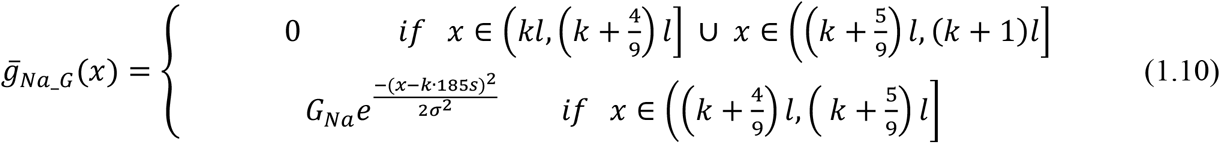

where *G*_*Na*_ = 300 *mS/cm*^2^, *k* = 1, 2, 3, …, and *l* = 185*s* is the distance between two consecutive actin rings.

## Results

To address whether the periodic arrangement of *Na*_*v*_ channels impacts action potential properties, we applied the H-H model to the action potential and compared the results generated by a constant vs. periodic maximum conductance using the rates and conditions described in Kole et al. [23-25]. To numerically solve the H-H model, we developed MATLAB-based code. For the case of a constant maximum conductance, which corresponds to homogeneously distributed *Na*_*v*_ channels, we made several observations: 1) The propagating action potential preserved its waveform as it travelled along the axon (illustrated in Fig. 2A,C). 2) The action potential peak amplitude was independent of the distance traveled; thus, the peak did not attenuate as it travelled across the axon. The difference between the resting membrane potential and the peak voltage of the action potential was 124 *mV*. The action potential propagated at a constant speed of 2.2 *m*/*s* (Fig. 2C). The period of the action potential was 1.2 *ms* and the wavelength (peak-to-peak distance) was 2.64 *mm*. Our numerical results are identical with the solution obtained by Kole et al. [24, 25] using the NEURON simulation environment. This confirms that our numerical method is able to accurately solve the H-H model and simulate an action potential and its propagation [23].

Next, we investigated the impact of distributing *Na*_*v*_channels in a periodic configuration matching what has been observed using super-resolution microscopy. Specifically, we randomly positioned *Na*_*V*_ channels in azimuthal bands repeated periodically along the axon. Each of these bands was positioned in the middle area between two consecutive actin rings and was approximately 20*s* wide (Fig. 1B, C). The random position of *Na*_*v*_ channels within the azimuthal strips followed a Gaussian distribution around the center line of each strip. The corresponding maximum conductance 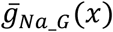 is given by Eq. 1.10. We also set the *Na*_*v*_ channel maximum conductance *G*_*Na*_ of the Gaussian distribution to be same as the maximum conductance *G*_*Na*_in a homogeneous distribution (300 *mS/cm*^2^).

Using this scenario, we ran the same simulations as we did for the homogenous distribution of *Na*_*v*_ channels. We found that the resulting action potential was different than the action potential generated by homogeneously distributed *Na*_*v*_ channels. For instance, the peak amplitude of the action potential was 19 mV, 6-7 times smaller than in the homogenous conditions, the propagation speed was 1.2 m/s (compared to 2.2 m/s), and the wavelength was 0.84 mm, 3 times smaller than the corresponding value in homogenously distributed *Na*_*v*_channels (Fig. S1).

We reasoned that the difference between the two scenarios was due to the lower number of *Na*_*v*_ channels per area (surface density) between two consecutive actin rings for periodically distributed *Na*_*v*_channels compared to homogeneously distributed *Na*_*v*_ channels. As mentioned above, periodically spaced *Na*_*v*_ channels are randomly positioned within an azimuthal band following a Gaussian distribution (see Eq. 1.10). Integrating the surface density of the *Na*_*v*_ channels, we found that the total number of *Na*_*v*_channels between two consecutive actin rings was approximately 60% of the homogeneously distributed *Na*_*v*_channels, when applying the same maximum conductance for simulations in the homogenous 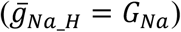 and periodic (Eq. 1.10) configurations.

Thus, we repeated our simulations with increasing maximum conductances in the Gaussian distribution to identify the maximum conductance value that produces an action potential similar to the action potentials derived under the homogenous conditions. We found that increasing the maximum conductance G_*Na*_ of *Na*_*v*_by 15 times reproduced the action potential properties of the homogenous configuration. The maximum conductance in this case is given by the expression:

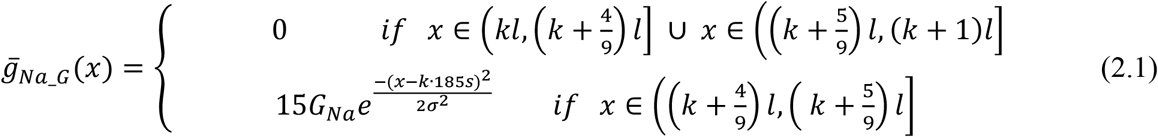

Thus, by solving the H-H equation using the maximum conductance shown in Eq. 2.1, we obtained the action potential illustrated in Fig. 2B, D. Under these conditions, the generated action potential had very similar properties (i.e., peak amplitude, period, wavelength, propagation speed) as the action potential generated by homogeneously distributed *Na*_*v*_ channels. The reason for this similarity is that, by increasing the *Na*_*v*_ maximum conductance to 15 *G*_*Na*_, we achieved the same number of *Na*_*v*_ channels between two consecutive actin rings as with the homogeneously distributed *Na*_*v*_ channels, for which 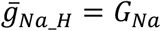. Thus, as long the number of *Na*_*v*_ channels is the same between the two consecutive actin rings, their distribution has no impact on the generation or propagation of the action potential. In retrospect, this is not surprising, as the wavelength of the action potential is on the order of 2.6 *mm*, which covers an area of approximately 1.4 × 10^4^ actin rings, meaning that the effect of the local distribution of *Na*_*v*_ channels is averaged over the entire wavelength.

Although the *Na*_*v*_ channel periodic distribution did not directly impact action potential generation or propagation, our model suggests substantial effects on the sodium action currents. This is best shown in Fig. 3, which plots the sodium currents across the axon as they drive the travelling action potential. First, the sodium current amplitude per unit area was much larger in the periodic configuration. This is because the surface density of *Na*_*v*_ channels in the nanostrips is 15 times greater in the periodic configuration than in the homogeneous configuration. Second, rather than following a smooth activation and deactivation curve across space, the sodium current followed a rapid fire-like behavior. We also plotted the sodium current as a function of time averaged across areas, corresponding to 5, 10, and 20 actin rings and centered on a 1.6 *mm* distance from the left end of the simulated axon (Fig. 4). This step is necessary, as our knowledge on sodium currents in neurons is through whole-cell recordings that record the activity across a large area. We found that the overall currents for different numbers of rings had very similar properties (i.e., peak amplitude, time course) independent of whether the *Na*_*v*_ channels were distributed homogenously or periodically. We found the same result whether we measured the current at 1.2 *mm*, 2.0 *mm*, or 2.4 *mm* from the left end of the axon. Thus, macroscopic measurement of sodium currents cannot distinguish on whether the channels are arranged in periodic or homogenous manner.

**Fig 4.**
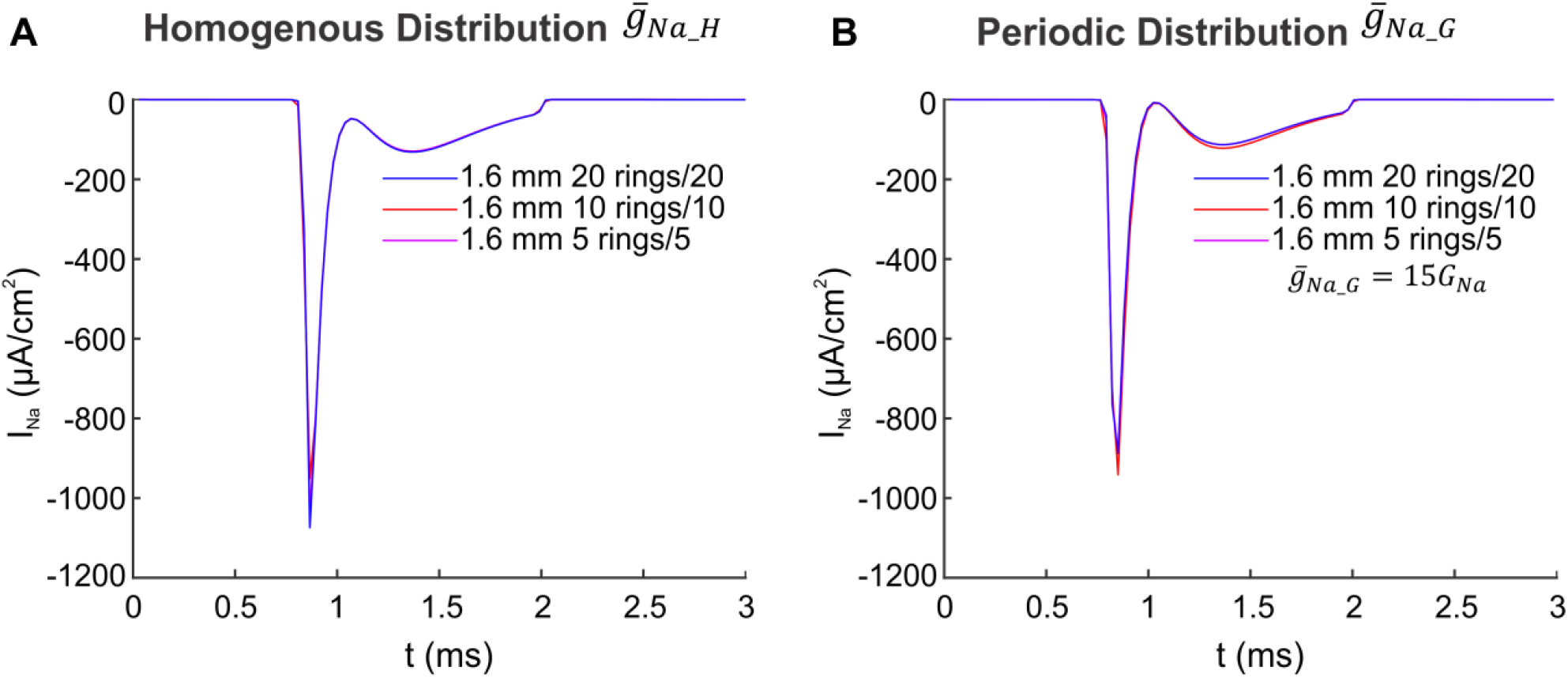
Illustration of the average sodium current as a function of time. Sodium current as a function of time averaged over areas corresponding to 5, 10, and 20 rings centered at a 1.6 *mm* from the left end of the axon for (A) homogeneous and (B) periodic distribution of *Na*_*v*_channels.

Overall, our simulations suggest that the periodic distribution of *Na*_*v*_channels primarily shifts the distribution of channels into “nano-strips,” leading to the emergence of *Na*_*v*_ channel nanodomains. These repeated nanodomains of axonal *Na*_*v*_ channel distribution do not affect the generation or propagation of the action potentials, but do alter the time course of the sodium currents locally. However, we predict that these changes can be measured not at a micrometer length scale but at the nanodomain level.

## Discussion

One of the major breakthroughs in our understanding of axon biology was the recognition that the actin-spectrin scaffold network in the axon follows a unique anatomical arrangement. In particular, super-resolution microscopy revealed that the plasma membrane skeleton has a periodic structure comprising of actin rings connected by spectrin tetramers [2]. These experiments also showed that *Na*_*V*_ channels exhibit a periodic ring-like distribution pattern that alternates with actin rings and co-localizes with ankyrin-G and ankyrin-B proteins [2, 3]. A coarse-grain molecular dynamics model of the axon membrane skeleton showed that the spectrin tetramers are under entropic tension and that the distance of a thermally moving ankyrin particle from its mean position follows a Gaussian distribution [5]. As a result, the ankyrin proteins and connected *Na*_*V*_ channels are distributed in azimuthal bands following a Gaussian distribution.

Although the role of the periodic actin ring structure in establishing the mechanical properties of the axon has been well established, it is unclear whether this structure also impacts the generation and propagation of the action potential along the axon. Our current understanding of the influence of *Na*_*v*_ channels on action potentials stems from the seminal work of Hodgkin and Huxley [14, 16, 17]. However, an assumption of the H-H model is that *Na*_*V*_ channels are distributed in a homogenous manner, which is now understood to be incorrect [2, 26]. Thus, in this study we tested whether the periodic distribution of the *Na*_*v*_ channels in the axon influences the generation and propagation of action potentials. Based on our simulations, we propose that: (i) *Na*_*V*_ channel periodicity does not affect action potential properties. A key parameter for an action potential is the number of *Na*_*v*_ channels per surface area between two consecutive actin rings rather than their spatial arrangement. (ii) Periodic *Na*_*v*_ channels lead to the formation of high surface density *Na*_*v*_ channel nanodomains. Such nanodomains might lead to a very large sodium ion concentration similar to the nanodomains seen with calcium channels in presynaptic terminals.

Our conclusion that action potentials are influenced by the total number of *Na*_*v*_ channels per surface area rather than their arrangement in the axonal compartment is not surprising in retrospect. Multiple studies have shown that *Na*_*v*_ channels located on the soma and even the dendrites of neurons can generate action potentials [27, 28]. In these compartments, the actin skeleton is not as rings connected by spectrin [26]. Additionally, recent work has shown that expressing the voltage-gated sodium channels along with leak potassium or voltage-gated potassium channels in oocytes can lead to the generation of action potentials [29]. Similar to our results, the key parameter is the number of channels per surface area expressed on the oocyte. Thus, based on our simulations and previously published research, we believe it is unlikely that the periodic *Na*_*v*_ channel distribution is critical for action potential generation or propagation.

Although the periodic *Na*_*v*_ arrangement does not contribute to the action potential, our results indicate that it greatly affects the *Na*_*v*_ channel surface density. In particular, the *Na*_*v*_ channel surface density must be 15 times higher than previous estimates that assumed *Na*_*v*_ homogeneity. The large surface density of *Na*_*v*_ channels in the nanostrips is required to equalize the average number of *Na*_*v*_ channels per surface area in both the periodic and homogeneous models of *Na*_*v*_ channel surface distribution. This density is necessary to generate the same action potential, which ultimately depends on the average sodium current density and the number of *Na*_*v*_ channels. If the unique arrangement of the *Na*_*v*_ channel in the axon does not contribute to the action potential, what might be its role? We considered a few possibilities. First, a sodium nanodomain might increase the likelihood of capturing sodium by Na-K ATPases. For instance, a large sodium flux is more likely to drive the occupancy of Na-K ATPase to a sodium-bound state followed by slow translocation to the outside. Considering the energetic cost of clearing sodium from the cytosol [30], increasing the likelihood of Na-K ATPase pumps to capture sodium will lead to more efficient transport. Similarly, a large sodium nanodomain would also increase the probability of activating axonal low affinity sodium-activated potassium channels following an action potential [31, 32]. Thus, the periodic sodium channel arrangement might be optimized for sodium signaling and reduction of the energetic costs associated with action potentials. Further studies and modeling are required to test these ideas and whether voltage-gated potassium channels overlap or are out-of-phase with sodium channels, as recently shown for KCNQ2 channels in axons [33].

## Conclusions

In this study, we simulated the action potential in a cylindrical compartment using the H-H model with a periodic *Na*_*V*_ channel spatial distribution, as has been experimentally described. We found that the *Na*_*v*_ channel arrangement does not impact the generation or propagation of the action potential, but rather leads to high-density sodium nanodomains.

## Supplementary material

**Fig S1.**
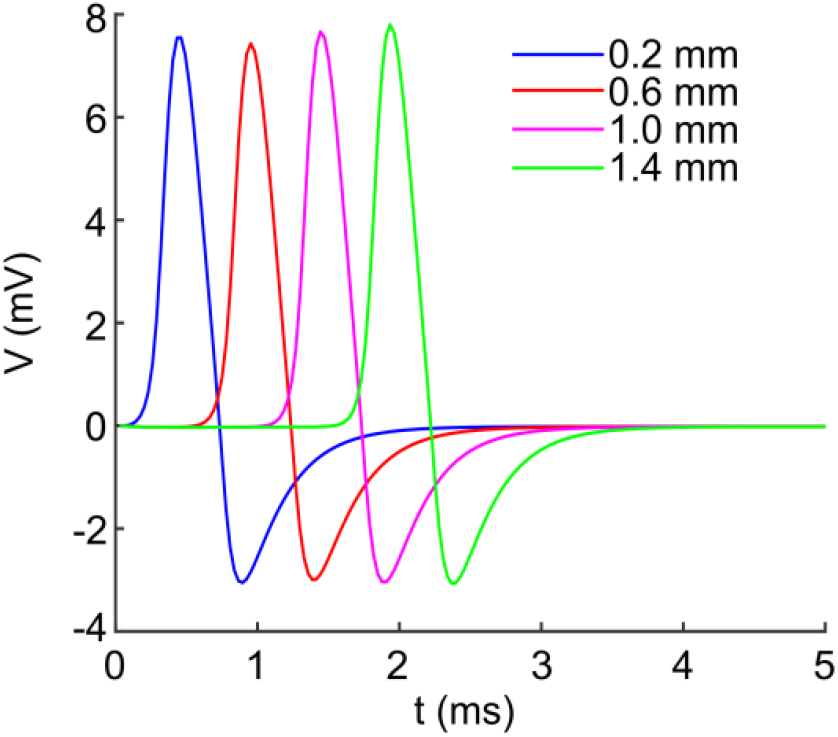
Illustration of action potential propagation for periodic *Na*_*v*_ channel distribution. The resting membrane potential was 0 *mV*. Illustration of action potential propagation for the case in which the *Na*_*v*_ channels are distributed periodically with a maximum conductance *G*_*Na*_ of the Gaussian distribution eq. 1.10 to be same as the *G*_*Na*_ under a homogeneous distribution configuration (300*m*S/c*m*^2^).

